# Diverse stimuli induce piloerection and yield varied autonomic responses in humans

**DOI:** 10.1101/2023.10.08.561417

**Authors:** Jonathon McPhetres

## Abstract

This research provides an in-depth exploration into the triggers and corresponding autonomic responses of piloerection, a phenomenon prevalent across various species. In non-human species, piloerection occurs in reaction to a variety of environmental changes, including social interactions and temperature shifts. However, its understanding in humans has been confined to emotional contexts. This is problematic because it reflects solely upon subjective experience rather than an objective response to the environment, and because, given our shared evolutionary paths, piloerection should function similarly in humans and other animals. We observed 1,198 piloerection episodes from eight participants while simultaneously recording multiple autonomic and body temperature indices, finding that piloerection in humans can indeed be elicited by thermal, tactile, and audio-visual stimuli. The data also revealed variations in cardiac reactivity measures: audio-visual piloerection was associated with greater sympathetic arousal, while tactile piloerection was linked to greater parasympathetic arousal. Despite prevailing notions of piloerection as a vestigial response in humans, it does respond to decreases in skin temperature and induces a rise in skin temperature during episodes. This research underscores that piloerection in humans is not solely an affective response to emotional stimuli. Rather, it is best understood as a reflexive response to environmental changes, suggesting a shared functional similarity with other species.

## Introduction

The integumentary system is a highly sensitive organ involved in many aspects of biological functioning, including in perceiving and responding to environmental changes. One of these responses is piloerection: the contraction of the arrector pili muscle, causing the hair to stand erect and forming visible bumps on the skin.

Evident across a diverse range of non-human species, piloerection can be triggered by a variety of stimuli. Avian species may ruffle feathers as part of courtship rituals^1^, to intimidate rivals^2^, or in response to cold to retain heat^3^. Among mammals, the phenomenon is noticeable during threats^4–6^ and mating displays^7^, and also as a thermoregulation mechanism^8^. This physiological process, therefore, plays a role in social interaction, signalling, and in maintaining the organism’s thermal equilibrium.

While piloerection serves clear functions in non-human animals, the same cannot be said with certainty when it comes to humans. Our comparative lack of hair relative to other mammals has led some to label piloerection as a vestigial trait^9–11^. As a result, its study in humans has predominantly focused on piloerection as an indicator of various emotions^12^ though the literature is absent of any evidence of this.

This discrepancy between models brings forth an intriguing question: given our shared evolutionary paths, is it not plausible that piloerection functions similarly in humans and other animals? Are we oversimplifying the human experience of piloerection by attributing it mostly to emotional experiences, overlooking the fact that humans can likely experience piloerection in response to a range of stimuli?

Adding to this complexity, various stimuli have distinct neural pathways. For instance, tactile stimuli are detected by afferent c-tactile fibres, and thermal information is passed from thermoreceptors along afferent Aδ fibres^13^. In contrast, audio-visual stimuli—be it a moving symphony or an unexpected sound and rustling in the treeline—utilize optical and auditory routes, and these engage a different set of neural processes. Yet, these diverse routes all converge to activate the same efferent sympathetic fibers^14^, which innervate the arrector pili muscles. This leads to a critical question: given that multiple paths converge to initiate the same final response, is piloerection a singular physiological reaction, or do the underlying autonomic responses reflect the diverse stimuli?

## Method

The study was designed to maximise the number of piloerection events observed within each participant. This research involved eight right-handed participants (k = 1,198 piloerection events), five females and three males, aged between 20 to 63 years old (M = 28.63, SD = 15.39) with BMIs ranging from 15.5 to 34.2 (M = 28.6, SD = 15.4), drawn from the university and surrounding community. Participants arrived in a laboratory wearing shorts and a t-shirt where they were connected to physiological equipment (MP160, BioPac System Inc.) to capture a variety of signals including a Lead II electrocardiogram, impedance cardiography, and continuous blood pressure. Sublingual and axillary thermometers provided a continuous record of core body temperature and axillary skin temperature, respectively. Physiological metrics were calculated using MindWare (Mindware Industries) pre-processing software. Piloerection events were observed through four high-definition cameras positioned to capture the dominant upper dorsolateral arm, dominant dorsal calf, and both left and right anterolateral thighs. Cameras were synced to Acqknowledge (v. 5.0) and viewed later in video coding software (BORIS v. 8^15^) second-by-second to identify the peak of each piloerection event.

The participants underwent an acclimation period and the final minute of this was used as the baseline reference period (see Supplementary Materials for additional details). After the baseline, they were exposed to a series of stimuli blocks intended to induce piloerection. The blocks were pseudo-randomized and the entire process spanned approximately three hours. For tactile stimulation, the participants were in a warm lab environment (M = 20.28°C, SD = .88°C). C-tactile afferents were stimulated by lightly brushing the participant’s skin with a feather and a metal “tickler” toy^16^, and administering two gentle puffs of air into the right ear using a rubber bulb aspirator^17^. These stimuli were applied to the area around each camera location and the nape of the neck.

Thermal stimulation targeted Aδ afferents by exposing participants to cold temperatures. This involved a 20-minute stay in a separate cold room (M = 16.43°C, SD = .41°C) while watching a neutral video, as well as direct skin contact with ice packs at each camera location and the nape of the neck, which was applied within the warm room and randomised in with the other stimuli.

In the audio-visual stimulation stage, participants were relocated to a private cubicle (within the warm-room) where they viewed a neutral calibration video, followed by four piloerection-inducing videos. Two videos were chosen from previous research^18^ and are reported to cause piloerection in about 60% of participants. Two additional videos were chosen to present a variety of content, one from a previous article^19^, and a fourth was selected by the author. Further details of the study design which are not relevant to this report are discussed in the supplementary materials.

## Results

In total, 1,198 piloerection episodes were recorded across the three blocks. Tactile, thermal, and audio-visual stimuli were equally likely to cause piloerection (*R*^2^ = .03, p = .180)—that is, all participants experienced piloerection in response to each type of stimuli, with the exception that one participant did not experience piloerection in the cold room. However, there was some heterogeneity in the number of piloerection episodes resulting from specific tasks within each block (η_p_^2^ = .10, *p* = .001), likely due to variation in task length (see Supplementary Materials). Thus, these results demonstrate that multiple stimuli routes yield a singular physiological response in human subjects.

### Audio-visual piloerection reflects sympathetic arousal

To examine autonomic arousal, cardiovascular metrics were calculated for the ten seconds around each piloerection event. A reactivity score for each metric was then computed by subtracting the baseline score from the piloerection epoch, and a linear mixed-effects model (with random intercepts for subject id) compared the three blocks on each reactivity score. A summary of key metrics of sympathetic and parasympathetic arousal indices is presented in Table 1, below. Full metrics are reported in Table S3.

**Table 1.** Key metrics of autonomic arousal during piloerection epochs compared between stimuli blocks.

| Indicators of Sympathetic Activation |  |  |  |  |
| --- | --- | --- | --- | --- |
|  | *Audio-visual*<br>MΔ | *Tactile*<br>MΔ | *Thermal*<br>MΔ | *Omnibus Variance* |
| Pre-ejection period | -6.35 | -3.95 | -3.44 | $\eta^2 = .034$, $p < .001$, |
| Heart Rate | 1.24 | -2.13 | -2.00 | $\eta^2 = .061$, $p < .001$, |
| Vascular Resistance | .27 | -2.63 | -1.52 | $\eta^2 = .046$, $p < .001$, |
| Cardiac Output | .13 | .59 | .17 | $\eta^2 = .043$, $p < .001$, |
| Respiration Rate | 1.06 | 2.04 | 1.82 | $\eta^2 = .029$, $p < .001$, |
| Indicators of Parasympathetic Activation |  |  |  |  |
|  | *Audio-visual*<br>MΔ | *Tactile*<br>MΔ | *Thermal*<br>MΔ | *Omnibus Variance* |
| Respiratory sinus arrhythmia* | .71 | -.24 | -.49 | $\eta_p^2 = .161$, $p < .001$, |
| RMSSD^ | 3.75 | 10.87 | 2.14 | $\eta_p^2 = .025$, $p < .001$, |
| pNN50^ | 2.65 | 5.51 | -1.78 | $\eta_p^2 = .032$, $p < .001$, |

*Note:* MΔ scores are reactivity scores indicating mean change from baseline; \*model controls for BMI and respiration; ^model controls for BMI.

In short, audio-visual piloerection showed a greater decrease in pre-ejection period (a key indicator of sympathetic arousal), coupled with an increase in heart rate, vascular resistance, and cardiac output. In contrast, tactile and thermal piloerection showed *decreases* in heart rate and vascular resistance. Together, this pattern of results indicates that piloerection elicited by audio-visual stimuli is associated with a stronger sympathetic arousal pattern compared to tactile and thermal piloerection.

At the same time, tactile stimuli showed stronger evidence of parasympathetic arousal as indicated by a large increase in two out of three metrics of heart-rate variability: the root mean square of successive differences (RMSSD) and the proportion of normal inter-beat intervals greater than 50ms (pNN50). A third metric, respiratory sinus arrhythmia, showed results in the opposite direction, but RSA and pre-ejection period can frequently correlate positively (Weissman & Mendes, 2021).

### Tactile piloerection reflects dermatome innervation

Recent studies have demonstrated that piloerection occurs consistently across the body^18^, contrary to prevalent beliefs that it is more common on the forearm. I examined this here, as well, by computing correlations between the number of piloerection episodes at each anatomical location. Findings indicated that tactile piloerection showed notably lower correlations compared to those caused by audio-visual and thermal stimuli (see Table 2), which I attribute to the unique innervation patterns within each dermatome.

**Table 2.** Correlations between the number of piloerection episodes at each anatomical location.

| | Audio-visual<br>Average $r = .69$ | | | Tactile<br>Average $r = .39$ | | | Thermal<br>Average $r = .81$ | | |
| --- | --- | --- | --- | --- | --- | --- | --- | --- | --- |
| Variable | 1 | 2 | 3 | 1 | 2 | 3 | 1 | 2 | 3 |
| 1. R. Arm | - |  |  | - |  |  | - |  |  |
| 2. Dom. Calf | .53* | - |  | 0.13 | - |  | .99* | - |  |
| 3. R. Thigh | .53* | .84* | - | 0.23 | .65* | - | .74* | .77* | - |
| 4. L. Thigh | .46* | .83* | .92* | 0.18 | .39* | .75* | .74* | .73* | .89* |

Note: \* $p < .01$

Specifically, the largest correlation under tactile stimulation appeared between the right and left thigh (*r* = .75), likely due to the shared innervation by the L3 spinal nerve. Similarly, the right calf, served by the L4 nerve, demonstrated a stronger correlation with the right thigh (*r* = .65) compared to the left (*r* = .39). Notably, the upper arm, innervated by the C5 spinal nerve, showed weak correlations with lower extremity piloerection. In contrast, audio-visual and thermal stimuli showed much stronger correlations across the anatomical locations. This is potentially due to the stimuli input route: audio-visual stimuli were processed through optical and auditory channels, while thermal stimuli impacted the entire body, as in the case of exposure to a cold room. Thus, these findings suggest that tactile piloerection is anatomically specific, triggered according to the location of sensory stimulation.

### Piloerection responds to, and affects, skin temperature

Contrary to the conception of piloerection as a vestigial trait, the data also showed that piloerection is associated with changes in skin temperature. To visualise this, 15 seconds of temperature data prior to and following the piloerection peak were selected. Because piloerection events across the body could occur within a few seconds of each other, each hypothesised to affect skin temperature, piloerection events were excluded if another event took place within that 15 second window. This allows for only prototypical events to be depicted. Finally, to test the hypothesis that more intense episodes of piloerection would result in greater changes to skin temperature, the intensity of the piloerection event was considered. Piloerection events were coded as “large” (fully formed bumps symmetrical around the infundibulum) or “small” (twitches or bumps in-between the infundibulum which dissipated before developing into large bumps).

Figure 1 shows that piloerection is preceded by a decrease in skin temperature, followed by an increase in temperature during the resulting piloerection epoch. The data were analysed in multiple ways. First, cross-lagged correlations (see Supplementary materials) indicated that the correlation between skin temperature was strongest about 8-10 seconds prior to the *peak* of the piloerection episode. Next, detrended axillary temperature at this lag time was used to predict the intensity of piloerection events (calculated as the rolling average of piloerection across all body locations) in a linear-mixed effects regression model. Larger decreases in skin temperature predicted more intense piloerection events (B = -.98, *p* < .001); oral temperature was not related to preceding temperature (B = .08, SE = .06, *p* = .150.

**Figure 1.**
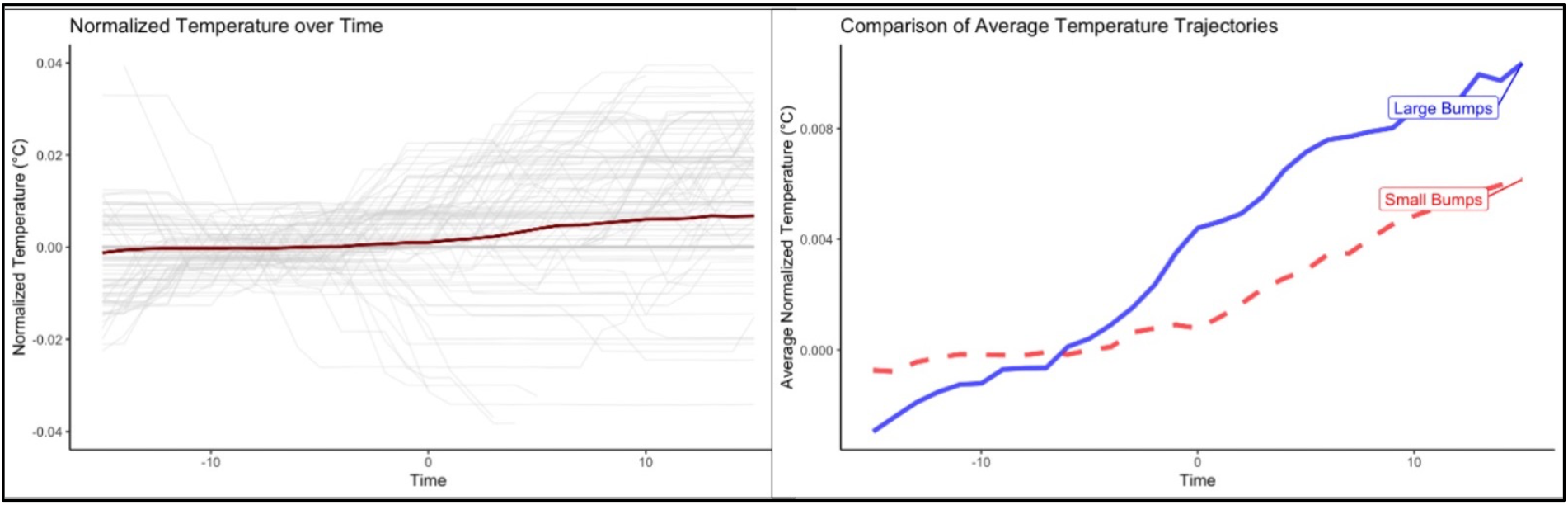
Piloerection is preceded by a decrease in skin temperature and followed by an increase in skin temperature during the piloerection epoch.

Second, skin temperature change from baseline around each piloerection event was predicted with dummy codes for task type in a mixed-effects model. Skin (axillary) temperature was higher during piloerection epochs, and this increase in temperature varied according to stimuli type (η^2 =^ .195, *p* < .001). Relative to baseline, skin temperature increased about .18°C (95% CI: -.02, .39) during audio-visual piloerection, .29°C (95% CI: .09, .50) during tactile piloerection, and .36°C (95% CI: .15, .57) during thermal piloerection. Core (sublingual) temperature, however, did not respond to stimuli type (η^2^ < .01, *p* = .108).

Finally, to test the hypothesis that more intense piloerection episodes would have greater effects on skin temperature, a linear mixed-effects model predicted skin temperature with Time and piloerection size. Results indicated that small piloerection events were associated with a continually smaller increase in temperature compared to large piloerection events (B = -.0002, *p* < .001). For example, at 15 seconds after the peak of piloerection, skin temperature during “large” piloerection events was about .004°C warmer than “small” piloerection events (Z = 6.33, *p* < .001, Cohen’s d = .62).

## Discussion

The prevailing understanding within existing literature treats piloerection in humans as a phenomenon separate from that observed in animal models. Discussion focused on animals treats piloerection as a response mechanism to a variety of stimuli, as a social signalling mechanism, and for thermoregulation, whereas humans are believed to experience this almost exclusively as an emotional response^12^. However, the findings from this study challenge this perspective. We found that piloerection in humans can be triggered by a diverse range of stimuli, including tactile, thermal, and audio-visual cues. These stimuli also had varying autonomic effects, including cardiovascular and thermal changes. These insights hold significant implications across the life sciences, as they underscore the necessity to review the dichotomous models traditionally used to interpret piloerection in humans versus animals.

One salient implication of our findings is that human piloerection can be induced by a similar spectrum of stimuli as those affecting animals. Although the functional role of piloerection in humans has become less pronounced—we no longer rely on hair erection as an indicator of impending threats, we have clothes for thermoregulation, etc—its physiological and sensory precursors have been conserved. Environmental changes bear a wealth of information and can spark significant physiological responses. These changes, whether in the form of acoustic and visual cues or temperature and tactile sensations, engender human reactions much as they would in animals. Thus, a gentle touch or a shift in temperature can induce piloerection, as could a sudden auditory or visual cue. The audio-visual stimuli employed in this study have much in common with the sensory experiences animals encounter in their natural environments. Moving music or frightening videos provoke in us physiological responses, akin to the reactions animals may display when confronted with distress signals from peers^5^ or potential threats^4^. This underscores a more comprehensive role of piloerection beyond subjective emotional responses in humans.

Our study further illustrates that these varied stimuli consistently led to changes in skin temperature. While core body temperature (as measured sublingually) remained unaltered, skin temperature demonstrated notable fluctuations prior to and during piloerection events. This highlights that despite previous characterisation of piloerection as a vestigial response in humans^10,20^, our findings offer substantial evidence that it retains elements of its functional characteristics.

The implications of our findings also have a far-reaching impact on the field of psychology. The primary interpretation of piloerection within psychological literature has been tied to emotional states. Our study, however, shows that piloerection should be reframed as a response to changes in environmental stimuli, shifting the focus away from its association with psychological experiences. Within this psychological literature, piloerection is commonly coupled with emotions typically linked to *parasympathetic* activity (for example, see a discussion around the emotion of awe^21–23^). However, it is well-established that the arrector pili muscles are innervated by the sympathetic nervous system^13,14^. Nevertheless, our study observed nuanced cardiovascular responses in conjunction with piloerection, contingent on the nature of the eliciting stimuli. For example, while audio-visual stimuli corresponded with elevated sympathetic cardiovascular activity, tactile stimuli led to heightened parasympathetic responses, which was found to be pleasurable in past resesarch^16^. These findings challenge the emotion-centric view of piloerection, which associates it with parasympathetic activation, and calls for a more comprehensive understanding of this physiological phenomenon in humans.

In conclusion, our study presents a transformative perspective on the occurrence and understanding of piloerection in humans. Contrary to the predominant view that assigns an almost exclusive emotional basis to this phenomenon, our findings suggest that piloerection is a nuanced physiological response to a variety of environmental stimuli. This aligns more closely with the multifaceted stimuli-response nature of piloerection observed in animal models. Furthermore, we demonstrate the need for a shift in focus from emotional to environmental stimuli within psychological literature on piloerection. The nuanced cardiovascular and thermoregulatory responses we observed, contingent on the nature of the stimuli, call for a more comprehensive and inclusive study of piloerection. Our research redefines piloerection as a complex and dynamic physiological response rather than a vestigial one, opening up new paths for future investigations in both physiological and psychological domains.

**Special thanks** to Samuel Forbes and Michael Lengieza for their help with R programming tasks for this project.

